# Comparability and Stability of Holotranscobalamin levels in Capillary and Venous Blood

**DOI:** 10.1101/2023.08.06.551133

**Authors:** Timothy Woolley, Emma Rutter, Macarena Staudenmaier

## Abstract

**Introduction:** Self-collected capillary blood has numerous advantages over venous samples taken via phlebotomy, as such the use of this finger-prick testing has increased over recent years, driven in some part by Covid antibody testing. However, there is limited evidence that venous and capillary serum is comparable for many routine analytes. In this study we aimed to determine whether capillary sampling could offer an alternative sampling method to venous for the assessment of vitamin B12 in the form of holotranscobalamin, and if this analyte was stable in unspun capillary blood for three days, thus allowing for samples to be returned by the postal system.

**Methods:** Matched pairs of capillary and venous blood samples were collected from 25 patients for the determination of holotranscobalamin, one set being processed on day zero, the other set being stored at ambient temperature and then processed on day three to mimic postal samples. Self-collected capillary blood was compared with phlebotomist-collected venous samples using correlation, Passing-Bablock regression, and Altman-Bland analysis.

**Results:** Passing-Bablock and Bland-Altman analysis showed holotranscobalamin levels in capillary and venous serum to be comparable and that the analyte was stable in unspun blood for at least three days at ambient temperatures.

**Conclusions:** We believe this is the first published study to determine if capillary blood sampling is an acceptable alternative to venous sampling for determining holotranscobalamin concentration; our data indicates that there is no significant difference in results from unspun venous and capillary blood stored at room temperature for at least 3 days compared to venous blood tested on the same day of collection.

## Introduction

Vitamin B12 is a dietary essential nutrient for humans, a deficiency of which can affect multiple systems including the bone marrow, where suppression leads to megaloblastic anaemia [1, 2]. The terminal ileum is the main site of Vitamin B12 absorption, where this process is aided by intrinsic factor (IF) produced in the gastric parietal cells. Causes of deficiency are numerous but include gastric atrophy, H. pylori, Crohn’s disease, drugs affecting absorption, and a strict vegan diet without corrective supplementation [1, 2, 3].

The resultant dyserythropoiesis leads to abnormal laboratory findings in several routine tests, including an elevated reticulocyte count, macrocytic red cells, and hyper-segmented neutrophils. Symptoms of deficiency typically include fatigue, visual disturbance, and memory loss. However, skin hyperpigmentation, glossitis, and infertility have also been reported [1, 2, 3, 4]

Within the United Kingdom, the prevalence of Vitamin B12 deficiency is approximately 6% in those less than 60 years of age and nearly 20% in those older than 60 years of age [1]. However, a population move to vegetarianism and veganism, could potentially increase these numbers. Routine use of screening tests in unselected patients is not recommended for the detection of Vitamin B12 deficiency but may be useful in specific situations [5, 6]. In these cases, a level of less than 150 pg/mL (111 pmol/L) is considered diagnostic for Vitamin B12 deficiency [1, 2, 3].

In serum, Vitamin B12 exists in 2 forms. It can be bound to haptocorrin to form holohaptocorrin or bound to transcobalamin to form holotranscobalamin (HoloTC). Cells can only take up Vitamin B12 in the form of HoloTC [7, 8].

Although not yet in widespread use, the measurement of HoloTC is however an emerging method of detecting deficiency, and there is growing evidence for it to be considered a better marker than Total B12 in the detection of early Vitamin B12 deficiency [9, 10]. Furthermore, HoloTC has been shown to be the best-performing indicator of Vitamin B12 status in the elderly population as well as during pregnancy, suggesting HoloTC may be suitable for screening of Vitamin B12 deficiency in these at-risk groups [9, 10, 11].

Routine Vitamin B12 and HoloTC status is typically performed on venous serum. However, conventional blood sampling using venipuncture is not always feasible or preferred by the patient. As such, alternative sampling strategies, including capillary sampling, are becoming commonplace.

Capillary sampling has several advantages over venous samples, which include being minimally invasive, requiring fewer raw materials, the ability to perform self-collection, and reducing wastage. Depending on analyte stability it also allows for blood to be taken at home and posted back to the laboratory.

In general, capillary samples consist of a peripheral blood sample obtained by a finger prick. It is assumed that capillary blood resembles venous blood, however, there are very few published reports on the comparison of analytes in capillary and venous blood. In addition, in many cases, capillary blood is still collected by a trained medical professional.

Using self-collected capillary samples opens several novel possibilities, including remote medical assessments and large-scale population screening. However, for the full potential of this method to be utilised, it is also imperative to assess analyte stability, thus allowing samples collected at home to be transferred to the laboratory using postal systems.

Our aim in this study was therefore to establish the validity of measuring HoloTC concentrations in capillary blood as a viable alternative to venous blood using two sets of samples, one pair (venous and capillary) being processed and tested on day zero, while the second pair (venous and capillary) would be stored at room temperature then processed and tested on day three. This second set of samples is being used to mimic samples being sent via the postal network.

## Materials and Methods

25 patients requiring HoloTC testing were asked to provide two venous and two capillary samples. Once consent was given, all participants were provided with diagrammatic guidance on how to perform effective capillary sampling, with self-collection performed immediately after the venous samples. Venous samples were collected using a trained phlebotomist. Venous samples were collected using Becton Dickinson (Oxford UK) Serum Separator Tubes (SST), while capillary samples were collected using Becton Dickinson Microtainer SST tubes and lancets. One pair of venous and capillary samples were then centrifuged together at 2000g for 10 minutes within 9 hours of collection and analysed together on the same day using the Roche (Burgess Hill, UK) Elecsys Active B12 assay on a Cobas e602 immunochemistry platform. The second set of samples were stored at room temperature (18-22 °C) for three days before centrifugation and again tested together on the same platform.

### Statistical analysis

Pearson correlation coefficient was used alongside Bland-Altman and Passing-Bablock to assess the strength of the relationship and level of agreement between HoloTC concentration in venous and capillary blood for samples taken on day zero and between venous serum from day zero and capillary serum from day three. All statistical analysis was performed using Analyse-It method validation edition (Analyse-It, Leeds, UK). Results Table 1 provides the absolute values for venous and capillary HoloTC concentrations (pmol/L) for each patient from both day zero and day three.

**Table 1.**
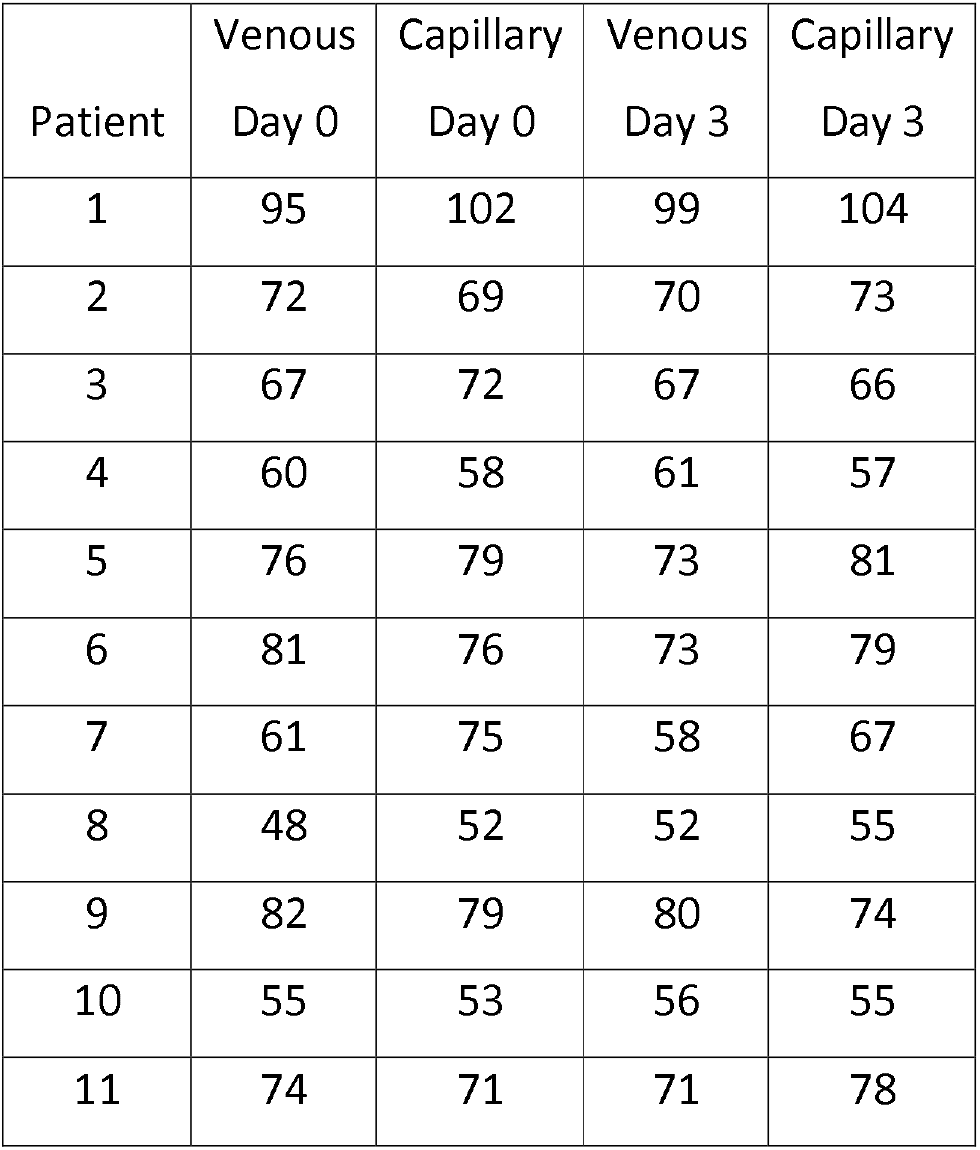
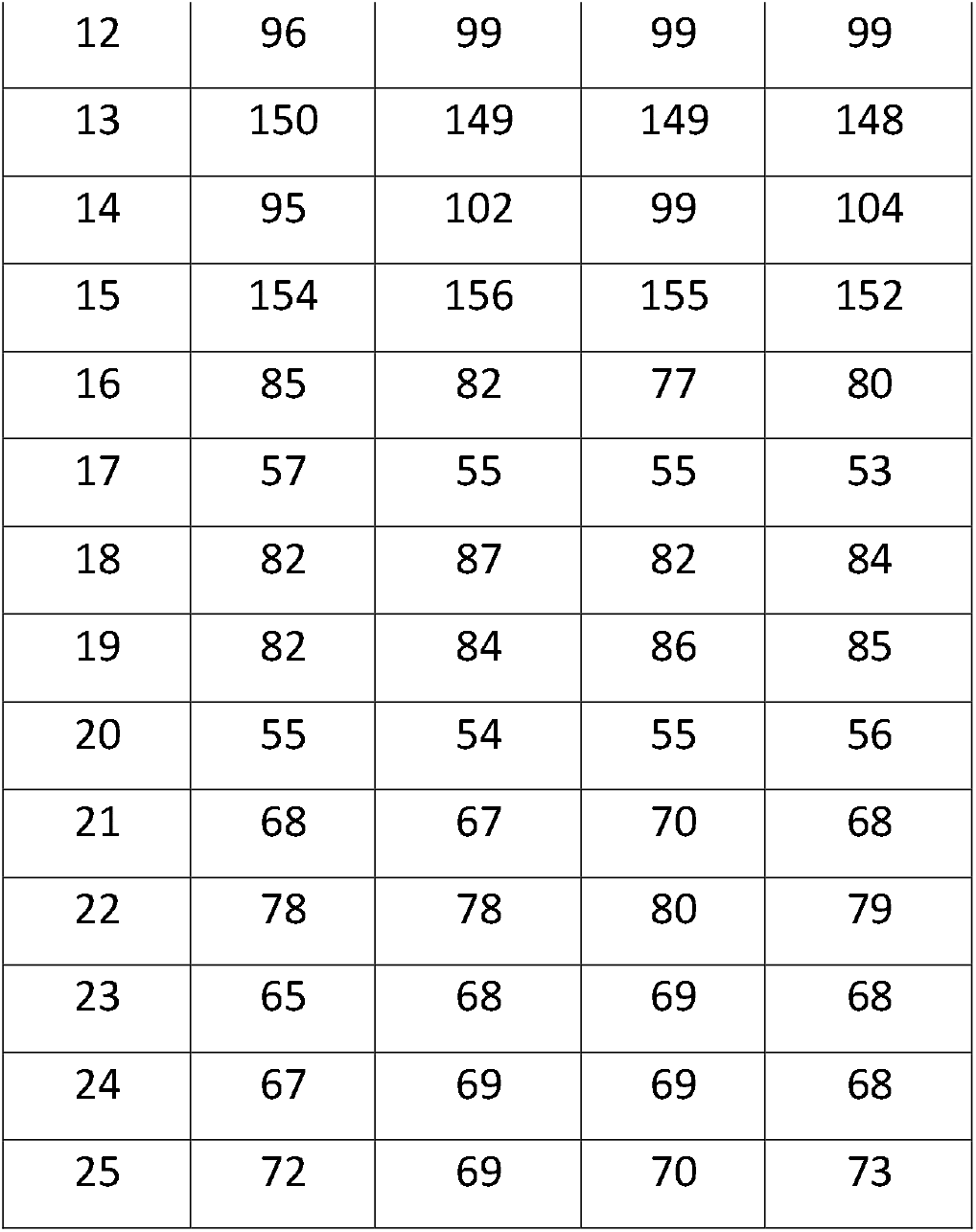
Day 0 and Day 3 - HoloTC Measurements in Venous and Capillary Serum (units pmol/L, n=25)

| Patient | Venous<br>Day 0 | Capillary<br>Day 0 | Venous<br>Day 3 | Capillary<br>Day 3 |
| --- | --- | --- | --- | --- |
| 1 | 95 | 102 | 99 | 104 |
| 2 | 72 | 69 | 70 | 73 |
| 3 | 67 | 72 | 67 | 66 |
| 4 | 60 | 58 | 61 | 57 |
| 5 | 76 | 79 | 73 | 81 |
| 6 | 81 | 76 | 73 | 79 |
| 7 | 61 | 75 | 58 | 67 |
| 8 | 48 | 52 | 52 | 55 |
| 9 | 82 | 79 | 80 | 74 |
| 10 | 55 | 53 | 56 | 55 |
| 11 | 74 | 71 | 71 | 78 |
| 12 | 96 | 99 | 99 | 99 |
| 13 | 150 | 149 | 149 | 148 |
| 14 | 95 | 102 | 99 | 104 |
| 15 | 154 | 156 | 155 | 152 |
| 16 | 85 | 82 | 77 | 80 |
| 17 | 57 | 55 | 55 | 53 |
| 18 | 82 | 87 | 82 | 84 |
| 19 | 82 | 84 | 86 | 85 |
| 20 | 55 | 54 | 55 | 56 |
| 21 | 68 | 67 | 70 | 68 |
| 22 | 78 | 78 | 80 | 79 |
| 23 | 65 | 68 | 69 | 68 |
| 24 | 67 | 69 | 69 | 68 |
| 25 | 72 | 69 | 70 | 73 |

**Figure 1.**
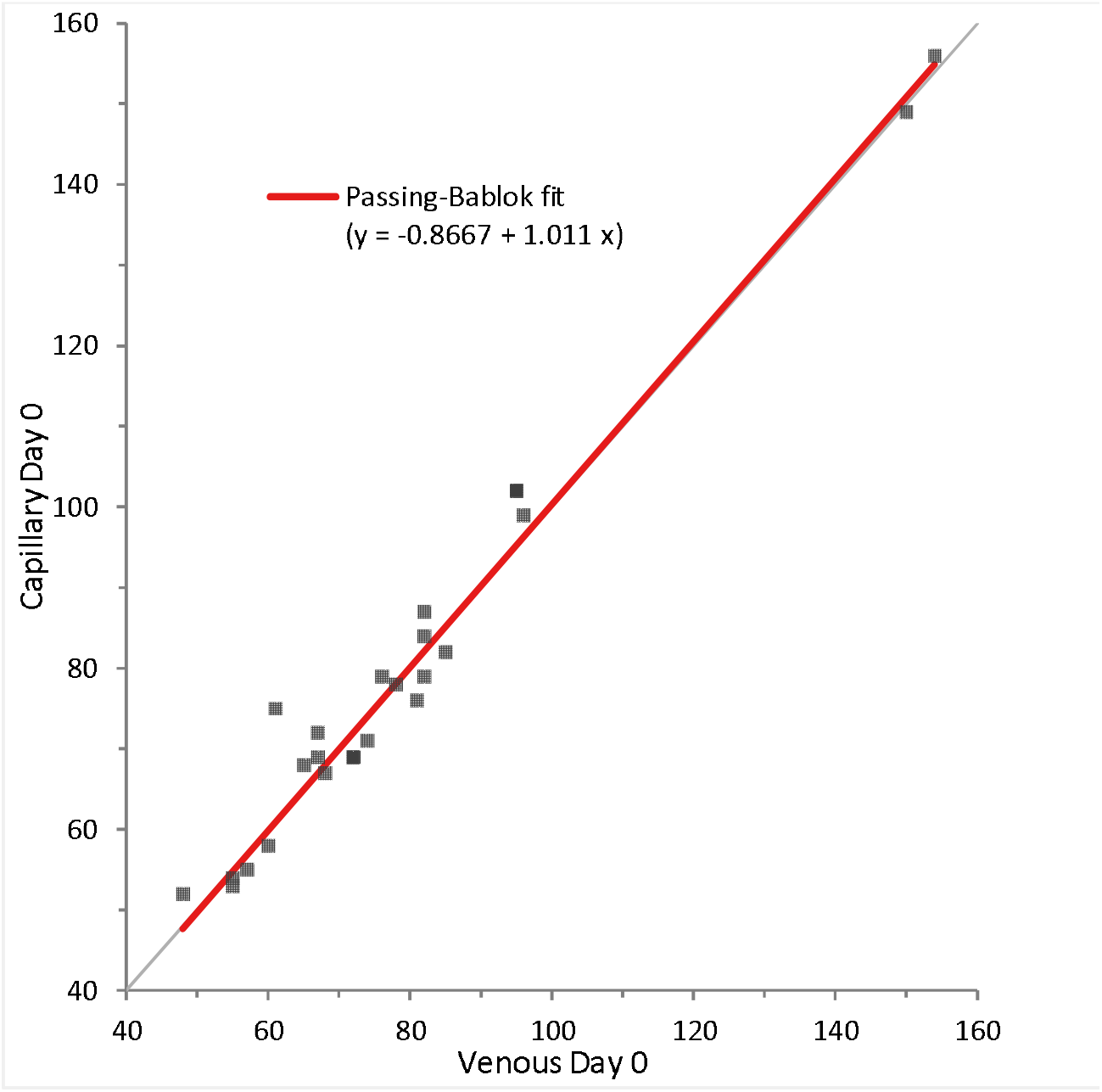
Passing-Bablock plot of venous and capillary serum determinations of HoloTC concentration on day zero (Correlation coefficient r=0.996, n=25) show a constant negative bias of 0.86pmol/L and a 1.1% positive proportional bias. No significant difference between venous and capillary serum is identified as the 95% CI encompass 1 for the slope and 0 for the intercept.

**Figure 2.**
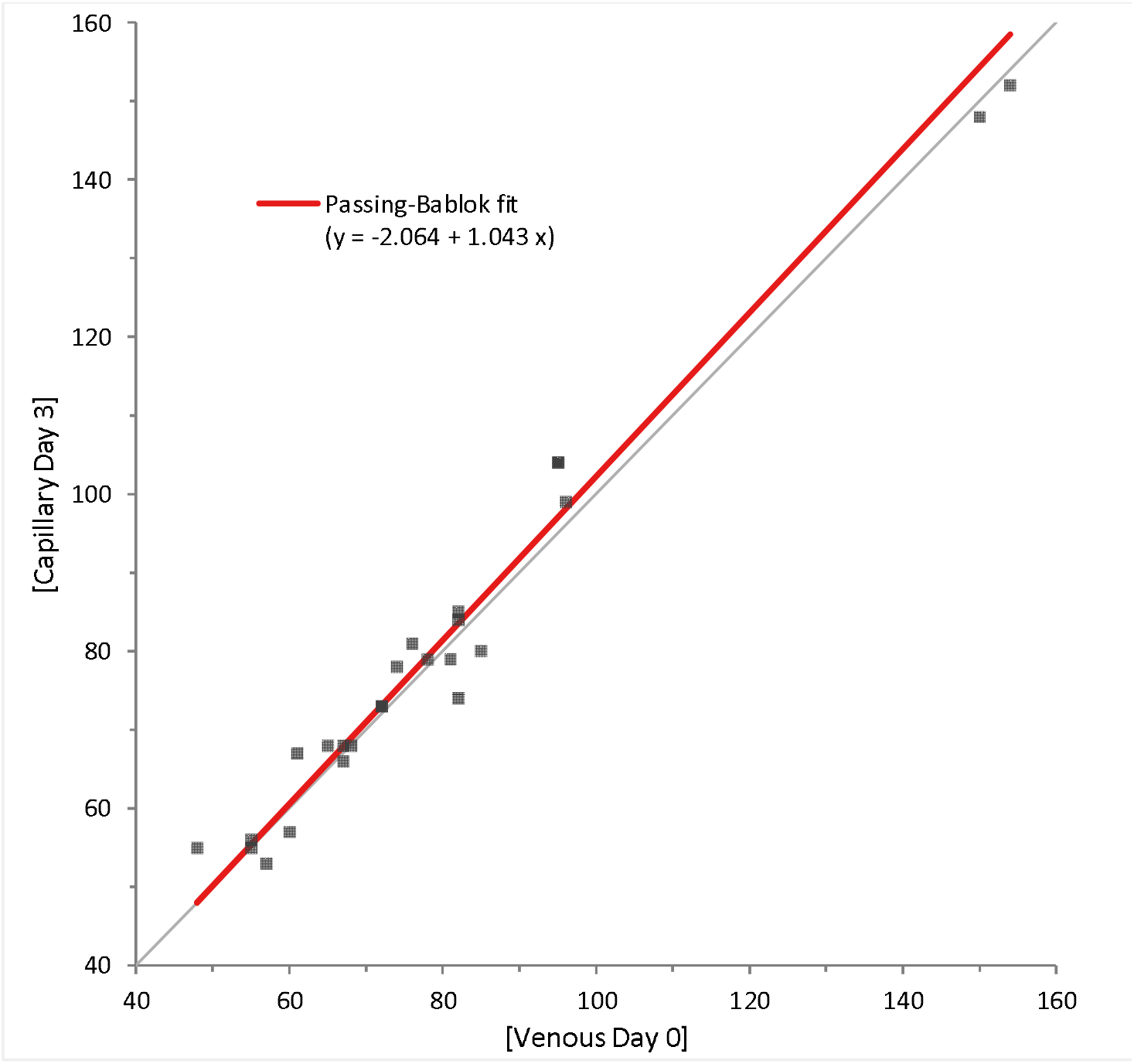
Passing-Bablock plot comparing venous HoloTC levels from day zero and capillary serum HoloTC levels on day three (Correlation coefficient r=0.987, n=25) shows a constant negative bias of 2.1pmol/L and a 4.3% positive proportional bias. No significant difference between venous Day 0 and capillary day 3 serum is identified as the 95% CI encompass 1 for the slope and 0 for the intercept.

**Figure 3.**
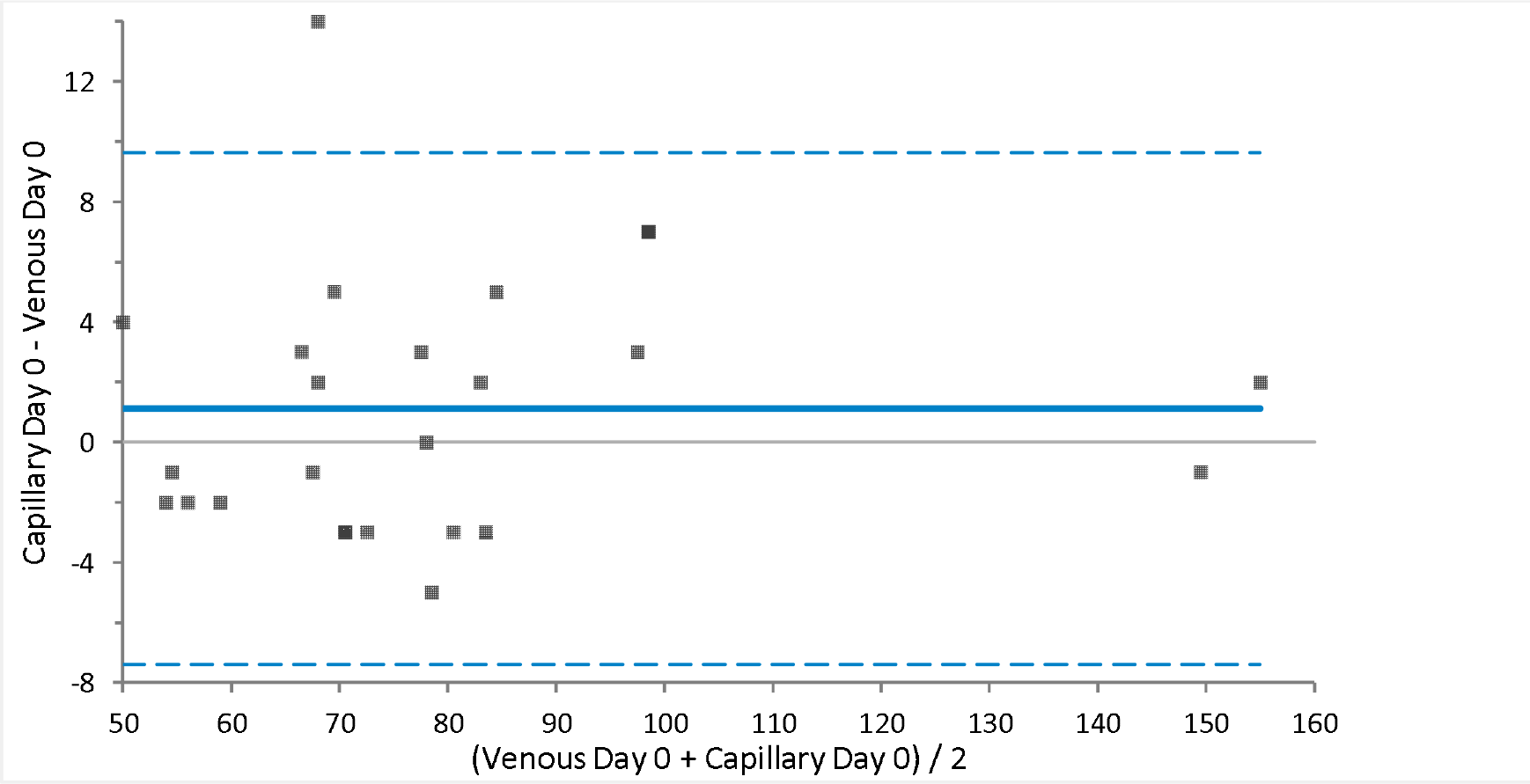
Bland-Altman plot of venous and capillary serum determinations of HoloTC concentration on day zero. Bland-Altman plot of agreement between venous and capillary (data from table 1, n=25) indicates a positive bias of 1.1 pmol/L. Limits of agreement are shown in blue dotted lines, line of identity is shown as solid blue line.

**Figure 4.**
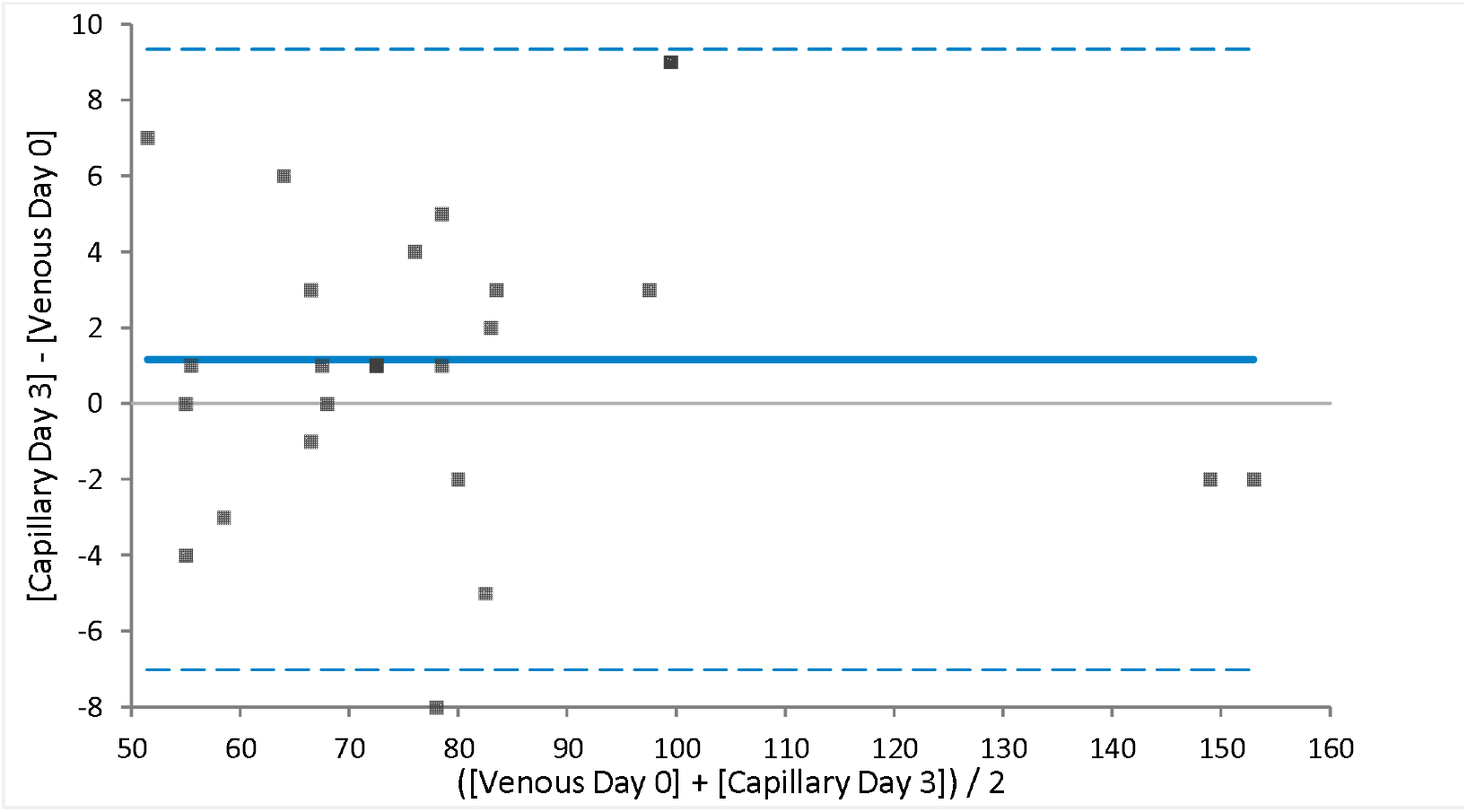
Bland-Altman plot comparing venous HoloTC levels from day zero and capillary serum HoloTC levels on day three indicates a positive bias of 1.2pmol/L. Limits of agreement are shown in blue dotted lines, line of identity is shown as solid blue line.

Our findings show that there is very good agreement between capillary and venous serum HoloTC concentration, both on day zero and on day three. A correlation coefficient of r = 0.986 and 0.987 (p<0.001) was observed between venous and capillary samples on day zero and venous serum from day zero and capillary serum from day three respectively. Bland-Altman analysis shows a good level of agreement with a small bias of 1 pmol/L between matrices.

## Discussion

There is growing evidence to support the use of HoloTC to diagnose early Vitamin B12 deficiency, especially in the elderly [9, 10, 11], as well as other potentially at-risk groups. Taking this into account, capillary blood sampling may present some advantages over traditional venous sampling, especially given capillary sample collection does not require attendance at primary or secondary care facilities.

Studies have shown capillary sampling to be preferred to venous collection in several situations [12, 13], in addition, this simpler and less-invasive technique also requires less raw materials and therefore produces less waste. Given the capillary sampling lends itself to self-collection and hence remote healthcare opportunities, any analyte selected must be stable for several days in the post. Although there are numerous papers validating the use of capillary blood for infectious disease testing [14, 15], at present there is limited published data on the comparability and stability of biochemical serum analytes in capillary and venous blood, and few manufacturers list capillary serum as an approved matrices in their instructions for use.

Our study is the first to compare capillary and venous samples for the determination of serum HoloTC concentration, whilst also providing whole blood stability data. Our findings show that there is very good agreement between capillary and venous serum HoloTC concentration, both on day zero and on day three. A correlation coefficient of r = 0.996 and 0.987 (p<0.001) was observed between venous and capillary samples on day zero and venous serum from day zero and capillary serum from day three respectively. In addition, Passing-Bablock and Bland-Altman analysis showed no clinically relevant bias between these matrices either on day zero or day three.

These results are of particular importance when investigating populations where there are concerns over the use of the more invasive venous sampling techniques or the cost and resources required for phlebotomy. This sampling method reduces the need for patients to visit primary care or outpatient clinics for blood tests, collection can be performed at home and thus could reduce non-attendance rates within the health service.

This method also lends itself to remote medical assessments, where samples can be taken at home and posted back to the laboratory for testing, especially as in this case the analyte of concern is stable for at least three days in whole blood at ambient temperatures. It is however critical that comparisons between blood matrices are included as part of the laboratories verification process as some analytes may behave differently between capillary and venous blood.

In conclusion, this work supports the use of capillary sampling for the determination of HoloTC concentration as an alternative to venous sampling, however it is noted that this study does not contain HoloTC concentrations at very low or high concentrations, likewise due to the difficulty is obtaining a series of controlled capillary samples, temperature stability testing was limited to ambient conditions.

The growth of self-collected capillary testing has made it critical for each laboratory offering this service to undertake a similar verification process during assay acceptance to confirm comparability between matrices. We hope that the evidence presented here will go some way to promote the use of capillary samples in routine pathology.

## Declarations

⍰Contributorship: All authors conceived the study. TW/ER were involved in data analysis. TW wrote the first draft of the manuscript. All authors reviewed and edited the manuscript and approved the final version of the manuscript.

⍰ Consent for publication - Consent for publication is given by the author(s)

⍰Availability of data and material - All data is provided within the manuscript

⍰ Competing interests - The author(s) declared no potential conflicts of interest with respect to the research, authorship, and/or publication of this article

⍰ Funding - The author(s) received no financial support for the research, authorship, and/or publication of this article.

⍰Ethics approval and consent to participate – Ethics approval was not required; all participants gave consent to participate.

## References

1. Hunt A, Harrington D, Robinson S. Vitamin B12 deficiency. BMJ. 2014 Sep 4;349:g5226. doi: 10.1136/bmj.g5226. PMID: 25189324

2. Stabler SP. Clinical practice. Vitamin B12 deficiency. N Engl J Med. 2013 Jan 10;368(2):149–60. doi: 10.1056/NEJMcp1113996. PMID: 23301732.

3. Dali-Youcef N, Andrès E. An update on cobalamin deficiency in adults. QJM. 2009 Jan;102(1):17–28. doi: 10.1093/qjmed/hcn138. Epub 2008 Nov 5. PMID: 18990719.

4. Toh BH, van Driel IR, Gleeson PA. Pernicious anemia. N Engl J Med. 1997 Nov 13;337(20):1441–8. doi: 10.1056/NEJM199711133372007. PMID: 9358143.

5. https://www.nice.org.uk/donotdo/tests-for-vitamin-b12-deficiency-should-not-be-carried-out-unless-a-full-blood-count-and-mean-cell-volume-show-a-macrocytosis. Accessed 23 August 2021

6. Devalia V, Hamilton MS, Molloy AM; British Committee for Standards in Haematology. Guidelines for the diagnosis and treatment of cobalamin and folate disorders. Br J Haematol. 2014 Aug;166(4):496–513. doi: 10.1111/bjh.12959. Epub 2014 Jun 18. PMID: 24942828.

7. O’Leary F, Samman S. Vitamin B12 in health and disease. Nutrients. 2010 Mar;2(3):299–316. doi: 10.3390/nu2030299. Epub 2010 Mar 5. PMID: 22254022; PMCID: PMC3257642.

8. Nexo E, Hoffmann-Lücke E. Holotranscobalamin, a marker of vitamin B-12 status: analytical aspects and clinical utility. Am J Clin Nutr. 2011 Jul;94(1):359S–365S. doi: 10.3945/ajcn.111.013458. Epub 2011 May 18. PMID: 21593496; PMCID: PMC3127504.

9. Bondu JD, Nellickal AJ, Jeyaseelan L, Geethanjali FS. Assessing Diagnostic Accuracy of Serum Holotranscobalamin (Active-B12) in Comparison with Other Markers of Vitamin B12 Deficiency. Indian J Clin Biochem. 2020 Jul;35(3):367–372. doi: 10.1007/s12291-019-00835-y. Epub 2019 Jun 12. PMID: 32647416; PMCID: PMC7326863.

10. Al Aisari F, Al-Hashmi H, Mula-Abed WA. Comparison between Serum Holotranscobalamin and Total Vitamin B12 as Indicators of Vitamin B12 Status. Oman Med J. 2010 Jan;25(1):9–12. doi: 10.5001/omj.2010.3. PMID: 22125690; PMCID: PMC3215392.

11. Valente E, Scott JM, Ueland PM, Cunningham C, Casey M, Molloy AM. Diagnostic accuracy of holotranscobalamin, methylmalonic acid, serum cobalamin, and other indicators of tissue vitamin B12 status in the elderly. Clin Chem. 2011 Jun;57(6):856–63. doi: 10.1373/clinchem.2010.158154. Epub 2011 Apr 11. PMID: 21482749.

12. Knoblauch MA, O’Connor DP, Clarke MS. Capillary and venous samples of total creatine kinase are similar after eccentric exercise. J Strength Cond Res. 2010 Dec;24(12):3471–5. PMID: 21132860.

13. Bajis S, Maher L, Treloar C, Hajarizadeh B, Lamoury FMJ, Mowat Y, Schulz M, Marshall AD, Cunningham EB, Cock V, Ezard N, Gorton C, Hayllar J, Smith J, Whelan M, Martinello M, Applegate TL, Dore GJ, Grebely J; LiveRLife Study Group. Acceptability and preferences of point-of-care finger-stick whole-blood and venepuncture hepatitis C virus testing among people who inject drugs in Australia. Int J Drug Policy. 2018 Nov;61:23–30. doi: 10.1016/j.drugpo.2018.08.011. Epub 2018 Oct 25. PMID: 30388566.

14. Bielen R, KocÖM Busschots D, Verrando R, Nevens F, Robaeys G. Validation of hepatitis C virus RNA detection using capillary blood by finger prick (GenXpert system)-Hepatitis C fingerprick study. J Viral Hepat. 2020 Jul;27(7):709–714. doi: 10.1111/jvh.13284. Epub 2020 Mar 13. PMID: 32106345.

15. Napierala Mavedzenge S, Davey C, Chirenje T, Mushati P, Mtetwa S, Dirawo J, Mudenge B, Phillips A, Cowan FM. Finger Prick Dried Blood Spots for HIV Viral Load Measurement in Field Conditions in Zimbabwe. PLoS One. 2015 May 22;10(5):e0126878. doi: 10.1371/journal.pone.0126878. PMID: 26001044; PMCID: PMC4441418.

